# Multilevel Development of Cognitive Abilities in an Artificial Neural Network

**DOI:** 10.1101/2022.01.24.477526

**Authors:** Konstantin Volzhenin, Jean-Pierre Changeux, Guillaume Dumas

## Abstract

Several neuronal mechanisms have been proposed to account for the formation of cognitive abilities through postnatal interactions with the physical and socio-cultural environment. Here, we introduce a three-level computational model of information processing and acquisition of cognitive abilities. We propose minimal architectural requirements to build these levels and how the parameters affect their performance and relationships. The first sensorimotor level handles local nonconscious processing, here during a visual classification task. The second level or cognitive level globally integrates the information from multiple local processors via long-ranged connections and synthesizes it in a global, but still nonconscious manner. The third and cognitively highest level handles the information globally and consciously. It is based on the Global Neuronal Workspace (GNW) theory and is referred to as conscious level. We use trace and delay conditioning tasks to, respectively, challenge the second and third levels. Results first highlight the necessity of epigenesis through selection and stabilization of synapses at both local and global scales to allow the network to solve the first two tasks. At the global scale, dopamine appears necessary to properly provide credit assignment despite the temporal delay between perception and reward. At the third level, the presence of interneurons becomes necessary to maintain a self-sustained representation within the GNW in the absence of sensory input. Finally, while balanced spontaneous intrinsic activity facilitates epigenesis at both local and global scales, the balanced excitatory-inhibitory ratio increases performance. Finally, we discuss the plausibility of the model in both neurodevelopmental and artificial intelligence terms.

## Introduction

Understanding the human brain remains the major challenge of biological sciences and has become the focus of considerable attention from both neurobiology and computational sciences (1–3). As a consequence of the recent success of neuro-inspired algorithms, including artificial neural networks and reinforcement learning, the fast-developing field of machine learning continues to look to neuroscience for inspiration. For instance, recent reflections on the connectomic implications of brain *Hominization* in the course of evolution and brain development have underlined the importance of a still under-evaluated notion of multilevel processing in the brain from sensory processing – and its elementary “local” neuronal circuits – to higher brain functions (4). Several recent theories emphasize the hierarchical relationship between local and global processes in the making of higher-level cognition. Among these, the Global Neuronal Workspace (GNW) (5) is exemplary. A hierarchical relationship between local and global processes is also outlined in Kahneman’s functional distinction of “System 1” handling fast and nonconscious cognitive processes, and “System 2” handling cognitive tasks requiring slower and more concerted conscious effort (6). While current algorithms in machine learning are now tackling System 1 tasks at the human performance level, a major milestone for modern artificial intelligence would be to provide models capable of approximating System 2 cognition (7).

To a large degree, the strides made in artificial intelligence research during recent decades can be explained by the overwhelming success of error backpropagation. Nevertheless, its biological plausibility remains a matter of considerable debate (8–10). Unlike artificial deep networks, and due to their discontinuous nature, spiking neurons are thought to learn by other means. Concretely, electrophysiological methods have indicated two principal mechanisms of learning in the human brain: Hebbian learning (11) and reinforcement learning (12). The mechanism of epigenesis by synapse selection (or synaptic pruning) is also known to play a significant role in both learning and biological development. Moreover, it has been demonstrated that neurogenesis can occur in adults (13) and that astrocytes may be implicated in synaptic modulation during learning (14). Despite solid evidence from joint anatomical, physiological and molecular investigations in the course of nervous system development, such mechanisms have been underexploited in brain modeling and computer sciences (15, 16). Recent work in neuroscience continues to uncover novel mechanisms at play during learning, which are far from being integrated into computer science. These and other mechanisms may provide important links, both within and between levels of organization, in diverse contexts – ranging from genes networks to long-range neuronal connectivity of the brain (17).

If detailed key mechanisms of learning in the brain have already been investigated, there is as yet no theoretical consensus on how these varied learning mechanisms interact in the brain (18–21). Yet it is known that the brain operates constantly through variation-selection mechanisms at multiple timescales (17) and thus supports the development of a multiscale and dynamical perspective on cognition and learning (22, 23). In artificial intelligence, new approaches have adopted such dynamical and multiscale aspects using attention (24) or social interaction between multiple agents (25). Basic developments inspired by neuroscience have also highlighted the importance of better capturing hierarchical relationships (7, 26, 27). Finally, the idea of biological learning without any inductive bias, typical of ANNs, has been criticized. On the contrary, biological investigations point dramatically to the embodied quality of cognition, perception, and action, as well the important role of embeddedness in natural and social milieus (28, 29).

In this paper, we present a framework for biologically plausible learning, using simulations of synaptic epigenesis that combine multiscale architecture with STDP and dopamine signaling. We hypothesize that synaptic epigenesis unfolds differently at local and global scales and that the requisite conditions for solving complex tasks associated with the global neuronal workspace include not only local but global epigenesis. We first identify necessary and sufficient conditions for proper learning of perceptual, cognitive, and conscious tasks. Next, we analyze how the identified factors influence performance in these tasks. After introducing the neurocomputational model, results are presented and the key role of dopamine and inhibitory neurons in the learning of higher cognitive tasks is discussed. Finally, we delineate future perspectives for the field of computational cognitive neuroscience and neuroAI.

**Model**

### 1. Overview of the Model and its neuronal components

We propose a computational model of an elaborate multilevel neural network able to pass cognitive tasks of increasing difficulty (Fig. 1). The task solving develops at three nested levels of hierarchical organization, where synaptic epigenesis may proceed at both local and global scales. The three different levels of structural organization show the increasing complexity of connectomic architecture with each nested level through a continuous progression. The first level, which we refer to as the “*sensorimotor level*”, deals with local sensory processing and classification of visual information: it requires local synaptic epigenesis. At the second level, or “*cognitive level*” (30), the network successfully passes a delay conditioning task: it mobilizes multiple cortical areas and their integration requires long-range axonal connections. The third level referred to as the “*conscious level*” is able to carry a trace conditioning task using a similar architecture as the *cognitive level* yet, with the addition of the necessary contribution of inhibitory interneurons.

**Figure 1.**
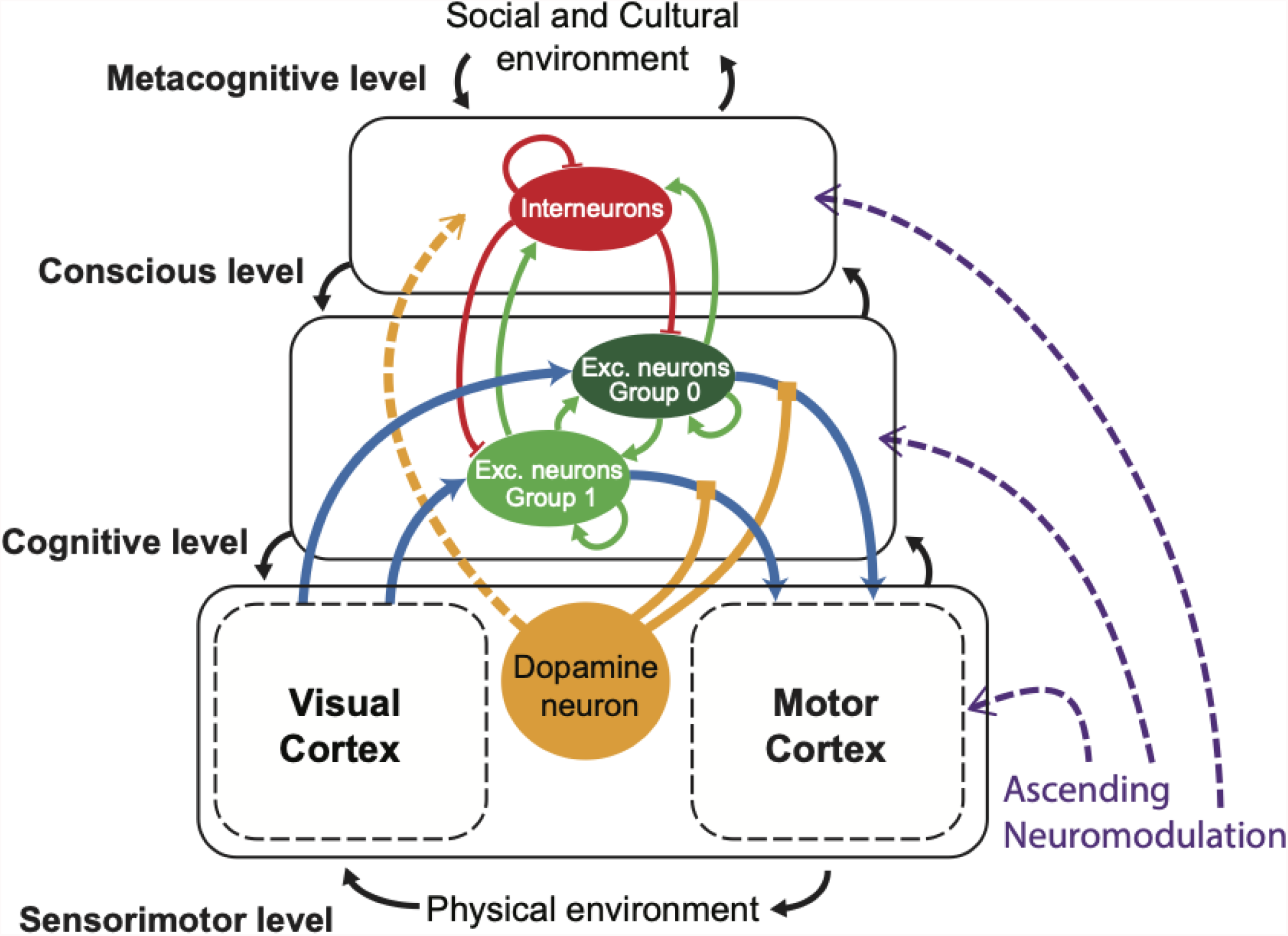
The full network architecture comprises Sensorimotor, Cognitive, Conscious, and Metacognitive levels. This is schematic representation of the full network and its components. Colors of arrows depict different types of connections: light blue - excitatory long-ranged (fixed: between VC and PFC and STDP: between PFC and MC), red - inhibitory (STDP), green - excitatory (STDP), yellow - extracellular dopamine modulation (affects dopamine modulated STDP). This version of the model is used to perform the trace conditioning task on the *conscious level*. To use the model with the delay conditioning task we do not introduce the interneurons population on the *cognitive level*.

Figure 1 illustrates the overall neuronal organization of the Model network. The three structural levels are all made up of neuronal layers organized hierarchically and nested within each other in a bottom-up and top-down manner (11). Each level may include several differentiated territories, or sub-networks, such as, for instance, the visual cortex or the motor cortex in the lower sensorimotor level. Every part of the network is modeled using neuronal assemblies except for the first two layers of the local network of the visual cortex. The state of each neuron is determined by the leaky integrate-and-fire model (31) and each assembly includes not only internal connections but external connections to other assemblies of the whole network. An important feature of the model is its evolution with time. In the course of its elaboration, the network connections are subject to epigenesis by selective stabilization of synapses (10). At critical stages of development, stable connections emerge from an initial growth process with overproduction of synaptic contacts with maximal diversity and variability, then a selection-stabilization regulated by the state of activity of the network takes place, together with the elimination – or pruning – of the un-selected ones (10). We examine, at all three levels of organization, how this development happens, both at local and global scales, and how internal and environmental factors affect its evolution.

The intrinsic spontaneous activity of the component neurons is a singular functional component introduced in the model proposed to occur at all its hierarchical levels. Also, the network is not exclusively composed of excitatory neurons as in most standard networks. At the highest level, we introduce inhibitory interneurons and test how the interneurons vs excitatory neurons ratio affects the performance. Long-range connections (5) which interconnect widely distinct territories of the global network are introduced at the higher cognitive and conscious levels.

Last throughout this work we use concomitantly different types of learning. At the local level, the network evolves under Hebbian learning with STDP (32). However, all the global connections are controlled by Reinforcement learning (12). To do so, we introduce the dopamine reward system (33), which is modulated by interactions with the external environment (Fig. 1).

### 2. Description of the tasks

The three selected tasks challenge differentially the sensorimotor, cognitive, and conscious levels.

The sensorimotor level is common to all tasks. At this level, we use the MNIST (Mixed National Institute of Standards and Technology) dataset of handwritten digits as visual input to the Visual Cortex (34). The network performs a *number recognition task* (35). Following image presentation, the network processes spikes arriving within the first 250ms, followed by a relaxation period of the same length. Thus, one image is shown every 500ms during the learning period. At the higher levels, when the full network receives an input from the Visual Cortex, the network is expected to respond to the question posed by the experimenter: “Is this number larger than x? If yes, push a button.”. For the case with only 0 and 1 in the input data, we took x equal to 0.

Two other tasks *delay* and *trace conditioning* (36), have been used to challenge the cognitive and conscious levels. In delay conditioning, the unconditioned stimulus (US) immediately follows or co-terminates with the conditioned stimulus (CS), whereas in trace conditioning, the CS and US are separated in time by a “trace” interval. Here, the numerical input operates as the Conditioned Stimulus (CS). We study two different variations of the task with different temporal locations of the CS and US. During the delay conditioning, we present a number for 150ms and the trigger stimulus coincides with the last 50ms of the input. To create the trace conditioning we first introduce an input for 150ms, followed by a pause of 200ms where the input is removed, and only after this pause the GNW receives the trigger stimulus of 50ms. Whenever the Motor Cortex produces output, i.e. the firing frequency of the motor neuron is above a certain threshold, the environment provides a negative or positive reward, depending on the answer.

### 3. Local network of the Visual Cortex

The architecture of this network (Fig. 2a), inspired by the work of Masquelier and Thorpe (37), consists of three layers representing the simplified model of the visual pathway. The first layer imitates the lateral geniculate nucleus and performs a convolution operation with filters of various angles. Then, in the second layer, only the strongest orientation reaches the V1 area. And in the last layer (V4 area) the patterns of digits are learned by mean of STDP. In the text we use the notation of ‘complex’ and ‘simple’ cells, thus the overall architecture can be written as S1-C1-S2 (see 37). A detailed description of each layer can be found in the Materials and Methods section.

**Figure 2.**
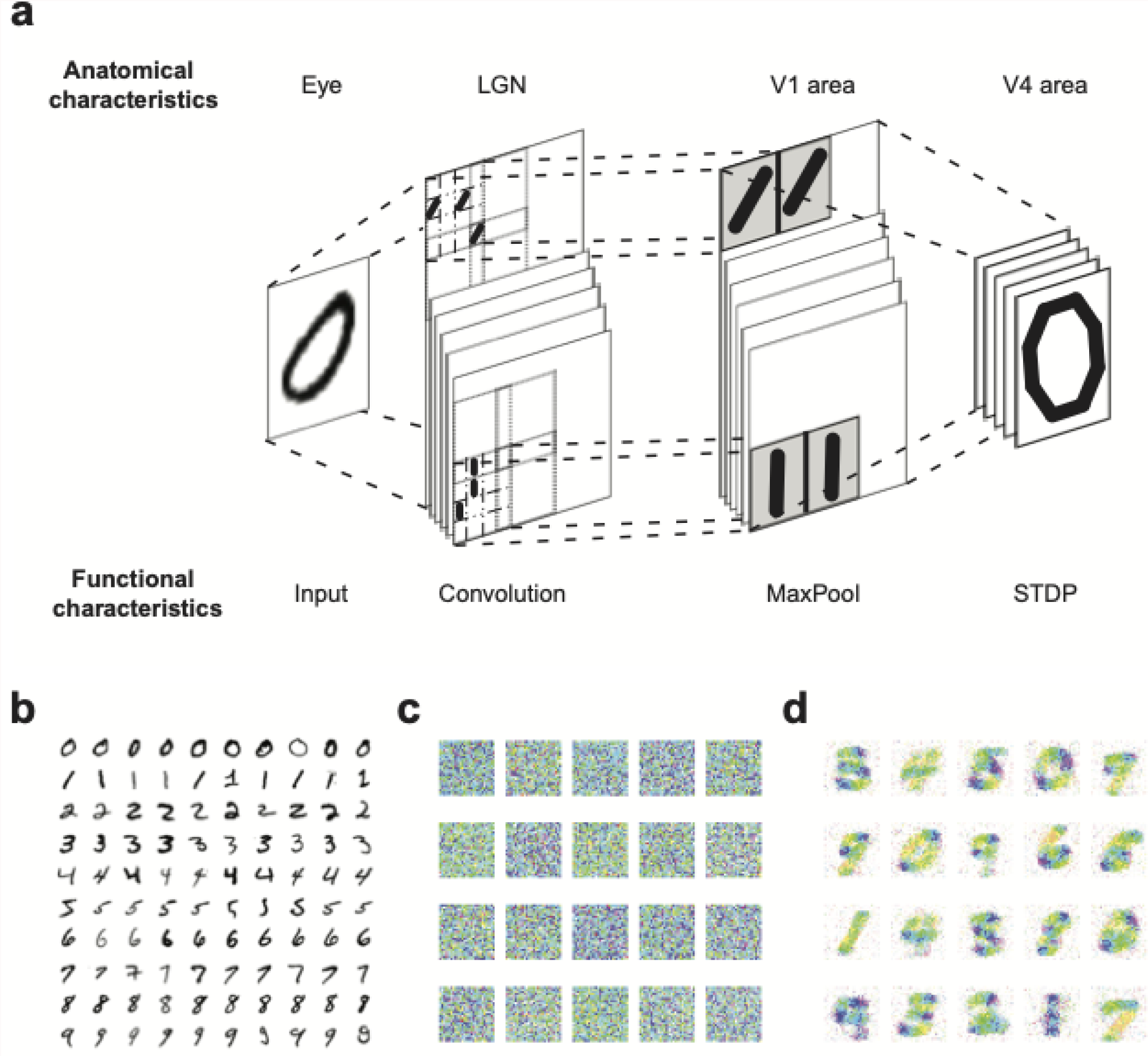
The local network of the Visual Cortex performs a classification task on the MNIST dataset. A. The schematic representation of the network structure with functional descriptions of each layer and corresponding anatomical analogies is shown. The model is based on three processing layers, where the first one performs convolutions on the original image with different filters of the same scale responsible for 6 different orientations. The second layer propagates only the strongest orientation in a given area. The connections of the last layer are regulated by STDP. The model is inspired by the paper of Masquelier and Thorpe, for more information see the original article (37). **B**. The sample of the MNIST dataset of handwritten digits, 100 examples. **C, D**. Pattern reconstructions of the 20 last layer cells before (**C**) and after (**D**) learning. The reconstructions were done using the HSV color scheme, where the color of each pixel represents the mean of the intensities throughout the 6 orientations at that location.

To train the model and observe the local epigenesis of STDP regulated synapses we used the MNIST dataset of handwritten digits (Fig. 2b). These inputs consist of images of handwritten digits from 0 to 9. MNIST is one of the fundamental datasets in Machine Learning, which makes it reliable and easy to use, as well as an informative benchmark for non-traditional learning methods. The images are introduced to the network continuously with equal relaxation periods in between. These relaxation phases are needed for the system to eliminate the temporal effect on STDP coming from the previous stimulus, and to bring all membrane potentials back to rest.

After a number of images are presented, the network’s plasticity allows it to evolve from a homogeneous state to one that has encoded various patterns by way of Hebbian learning. Fig. 2c-d shows what the patterns of 20 S2 cells look like before and after the network completes its learning process, respectively. These images are plotted using the HSV (hue, saturation, value) color scheme. Values for each pixel are set to 100%. Hue and saturation are determined in the following way: each pixel combines information from 6 synapses (by the number of used orientations). These 6 synaptic weights are transformed into the complex plane: from w to w* = w * exp(icp), where cp is the particular orientation. Next, all 6 weights are added together. The absolute value of the resulting complex number is saturation, and the angle is hue. Using this color scheme, we are able to see not only the strength of the C1-S2 connection but also the dominant orientation (if present) coming from any of the C1 cells (see bottom of Fig. 2d).

### 4. Full Network

After the local network has undergone the process of Hebbian learning, the weights of C1-S2 connections are frozen and this network is considered mature. Then, a new architecture is constructed on top of it, to explore global connections. This intermediate network consists of four fundamental elements: the Visual, Motor, and Prefrontal Cortices, as well as the Striatum, represented by a single “dopamine” neuron (Fig. 1). The Visual Cortex is essentially represented by the aforementioned local network with deactivated STDP and fixed synaptic weights. Input to the global network consists of the output of S2, once its trained model has extracted from the stream of images, features to be read by the full network. In other words, raw data is inaccessible to the full network, and only intermediate features can be accessed by subsequent brain regions. S2 output is passed to the excitatory population in the PFC.

The Motor Cortex is minimally represented by only a single neuron. The firing of this neuron is interpreted as revealing the interaction with the external environment such as pushing a button and giving a positive answer to a question. It processes outputs only from the excitatory populations in the PFC.

The Prefrontal Cortex is assumed to contain 2 populations of neurons: excitatory neurons and interneurons. To test the cognitive level with the delay conditioning, we use the simplest version of PFC without inter-neurons. The excitatory population is divided into 2 selective groups (by the number of digits we want to use for the task, in our case, it is a binary classification and we only use digits 0 and 1). Each selective group is connected to S2 cells in the Visual Cortex model which are responsible for one specific digit. All these connections are fixed. The connections inside the PFC between both excitatory and interneurons are all-to-all.

Learning in the PFC-MC connections is accomplished using reinforcement via interaction with the external environment (dopamine modulated STDP). The internal PFC connections are subject to Hebbian learning (classical STDP) in the same way as C1-S2 connections in the local network. More detailed information about STDP models can be found in the Materials and Methods section.

## Tasks solving by the Model

The proposed model of an elaborate multilevel neural network (fig1) is able to pass behavioral tasks of increasing difficulty. The three levels of structural organization exhibit increasing connectomic complexity with each nested level.

### 1. Sensorimotor level - visual recognition task

#### Stabilization and pruning

At this level, the neuronal network – described above as the “visual cortex local network” – can perform a recognition task of numbers, using handwritten digits as inputs. This image processing task mobilizes nonconscious perception, by leveraging local synaptic epigenesis as well as the spontaneous activity of the network. Fig. 3 a-d shows the computed multistep evolution of this performance.

**Figure 3.**
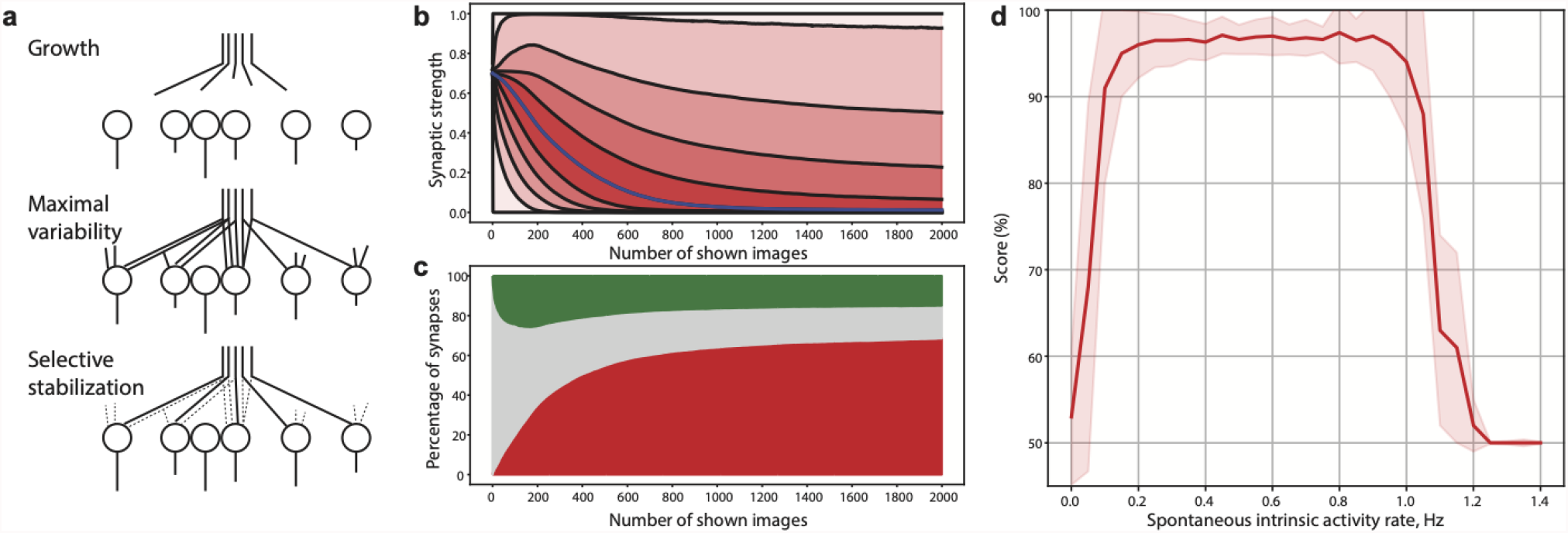
Local Epigenesis and performance of the Local Network. **a**. Schematic representation of the hypothesis of epigenesis by selective stabilization of synapses (from Changeux 1985 (38)) **b**. Evolution of the synaptic strength of the whole synaptic population in the last layer of the Local Network. Blue line: the median, black lines: 0-100th percentiles (with the step of 10). **c**. The epigenesis process in terms of selective stabilization of synapses, i.e. synaptic numbers instead of strength like in panel a. Red group: represents eliminated synapses with weights of less than 25%. Green group: represents selected synapses with weights of more than 75%. Grey group: includes weights between 25% and 75%. **d**. The performance of the matured network depends on the rate of spontaneous intrinsic activity. There is only a specific window where the network can produce a high performance of more than 90% with exact rates of the spontaneous intrinsic activity depending on the number of neurons.

The initial state of the LGN-V1 synaptic population is highly homogeneous. The synaptic weights are normally distributed around the mean of 70% with the standard deviation of 2%. When the network is presented with the constant input stream of images the synaptic population starts to evolve. The evolution of the synaptic strength of the whole synaptic population is presented in Fig. 3b. This graph shows the evolution of the percentiles for the population with a step size of 10, from 0 to 100. It can be seen that all synapses do not behave in the same way. Around two-thirds of population synapses immediately weaken and are as a consequence eliminated. At the same time, other synapses are strengthened and selected (see 10).

Fig. 3c describes the evolution of the population of synapses in terms of selective stabilization. The selected synapses are those responsible for the transfer of the signals related to the input information. Others are gradually eliminated, such that they exert a lesser impact on the final layer. However, this process is not linear and learning patterns change over time. Fig. 3d represents the evolution of the states within the synaptic population. In this schematic representation, a synapse belongs to the green (“selected”) area if the synaptic strength (or synaptic weight) is greater than 75%. Similarly, the red (“eliminated”) area accounts for synapses with a strength of less than 25%. All others between 25% and 50% fall into the gray (“undetermined”) area. This zone contains the entire population at the beginning of the simulation.

Due to non-linear behavior, we have outlined two distinct stages of learning depending on the growth rate: 1. from the start to ∼200 shown images and 2. from ∼200 to 2000. Statistical tests have been performed to establish that the growth rate of selected and eliminated synapses is reliably different during the first and second periods. The T-test has shown the p-value < 10^−8^ and Cohen’s D > 15 for both groups. With the chosen set of parameters, the first period occurs while the model is processing the first ∼200 images. During this time, we notice the rapid selective stabilization of half the synaptic population. Simultaneously the percentage of selected synapses rises either. During this period V1 cells gain their selectivity towards specific sets of stimuli.

The second stage of learning can be characterized as the fixation and reinforcement of the changes that happened before. The fraction of eliminated synapses monotonously reaches a plateau level with a value of around two-thirds of all synapses. The group of selected synapses, however, experiences “overshooting” and, after 200 images, starts to slowly decrease before reaching a plateau. This behavior can be described as a period when the S2 features become clearer, eliminating redundancy between them and within the spontaneous intrinsic activity.

In addition to synaptic epigenesis, the intensity of spontaneous intrinsic activity is among the factors affecting the performance of the network. In our model, each of the C1 neurons is able not only to produce spikes caused by the S1 layer but also to generate spontaneous intrinsic activity in a form of random Poisson distribution with a fixed rate. We can see that high performance (with the accuracy of classification > 90%) can only be achieved in a specific range of values (Fig. 3d). We observe no or very weak classification with values of spontaneous intrinsic activity close to zero; the same behavior is spotted for large values as well (> 1.0 Hz). The classification reaches high scores of >90% only between 0.1 Hz and 1.0 Hz. Thus, a sufficient but not exceedingly high amount of spontaneous intrinsic activity is a necessary factor in achieving the accurate recognition of numbers. The specific values of the spontaneous activity vary depending on how many neurons are used and how strongly it affects the conductivity of each neuron.

### 2. Cognitive level - nonconscious delay conditioning task

#### Long-ranged connections and reward

In this section, we examine which network global architecture is needed for the performance of a delay conditioning task (Fig. 2) again using digits as visual stimuli for the unconditioned and conditioned stimuli. We discover that both the presence of long-range connections and their epigenesis are required. The initial state of the network is homogeneous: the local connections are set to 1% of the maximum weight and the long-range connections between the Prefrontal Cortex and the Motor Cortex are 50% for all synapses (Fig. 4).

**Figure 4.**
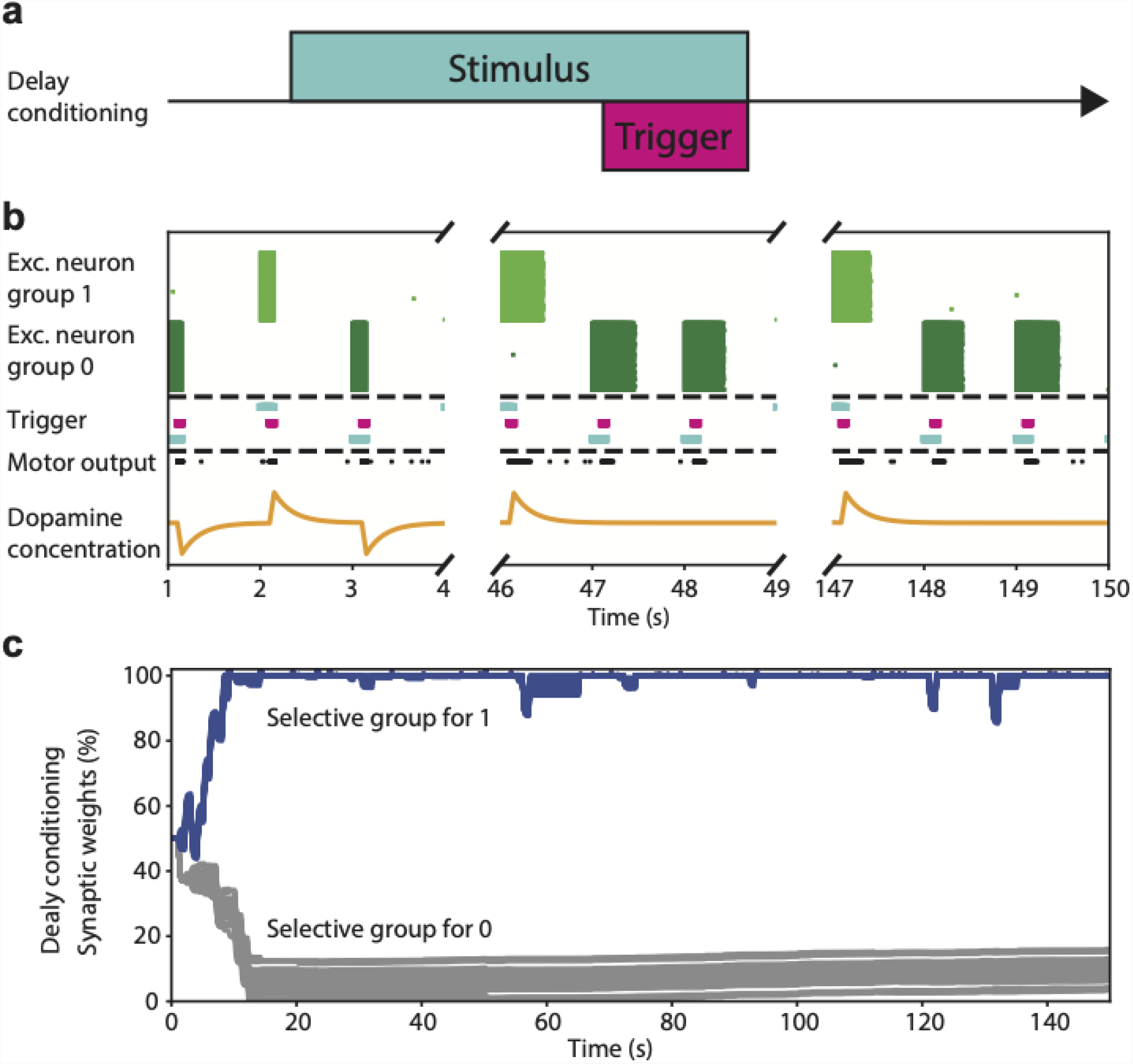
Epigenesis of the long-range connections network between Prefrontal and Motor Cortices while performing binary classification under the delay conditioning. **a**. Description of the delay conditioning task. b. Evolution of the firing rate of the two populations of neurons within the GNW (excitatory selective for the digit 1, excitatory selective for the digit 0). 15 neurons per population are shown, each dot shows a neuron spiking. Depending on the network parameters, if the internal excitation in the GNW is not strong enough to cause constant feedback firing, the stimulus representation exists only while the input signal is present. c. Epigenesis of the global connections between Prefrontal and Motor Cortices. The network needs less than 20 images to learn the task and makes mistakes only when the visual classification in the Local Network is wrong. Blue and grey lines correspond respectively to 1 and 0 output of the PFC.

The network uses long-range connections to transmit stimuli between Visual and Motor Cortices via a hub of excitatory neurons, as hypothesized in the Global Neuronal Workspace (GNW) model (5). We show that the network can perform a delay conditioned task (Fig. 4a represents the temporal relation between the stimulus and the trigger) without conscious processing. Fig. 4b depicts the activity of the Prefrontal Cortex at the beginning and the final stages of the learning process. The stimulus is erased from memory as soon as the input is gone, meaning that it is not processed in any way between Visual and Motor Cortices. However, depending on the initial parameters, our network can start firing constantly in a positive feedback loop since it contains only excitatory neurons. In this case, the network is not able to hold representations either. The epigenesis process transpires rather quickly, on the scale of ∼15 shown images, and all the synapses responsible for the representations of the digit 1 get selected while the representations of the digit 0 get eliminated (Fig. 4c).

The model turns out to be sensitive to the rate of spontaneous intrinsic activity. We have observed that, regardless of configuration, the model can learn how to perform the task only when the rate is set within a certain interval (Fig. 5). Moreover, while there is a specific range of values within which the network provides decent classification (accuracy > 95%), excessive activity can disturb the learning process, and the accuracy will remain around the chance level. The performance does not change significantly as long as the spontaneous intrinsic activity is set in the right interval.

**Figure 5.**
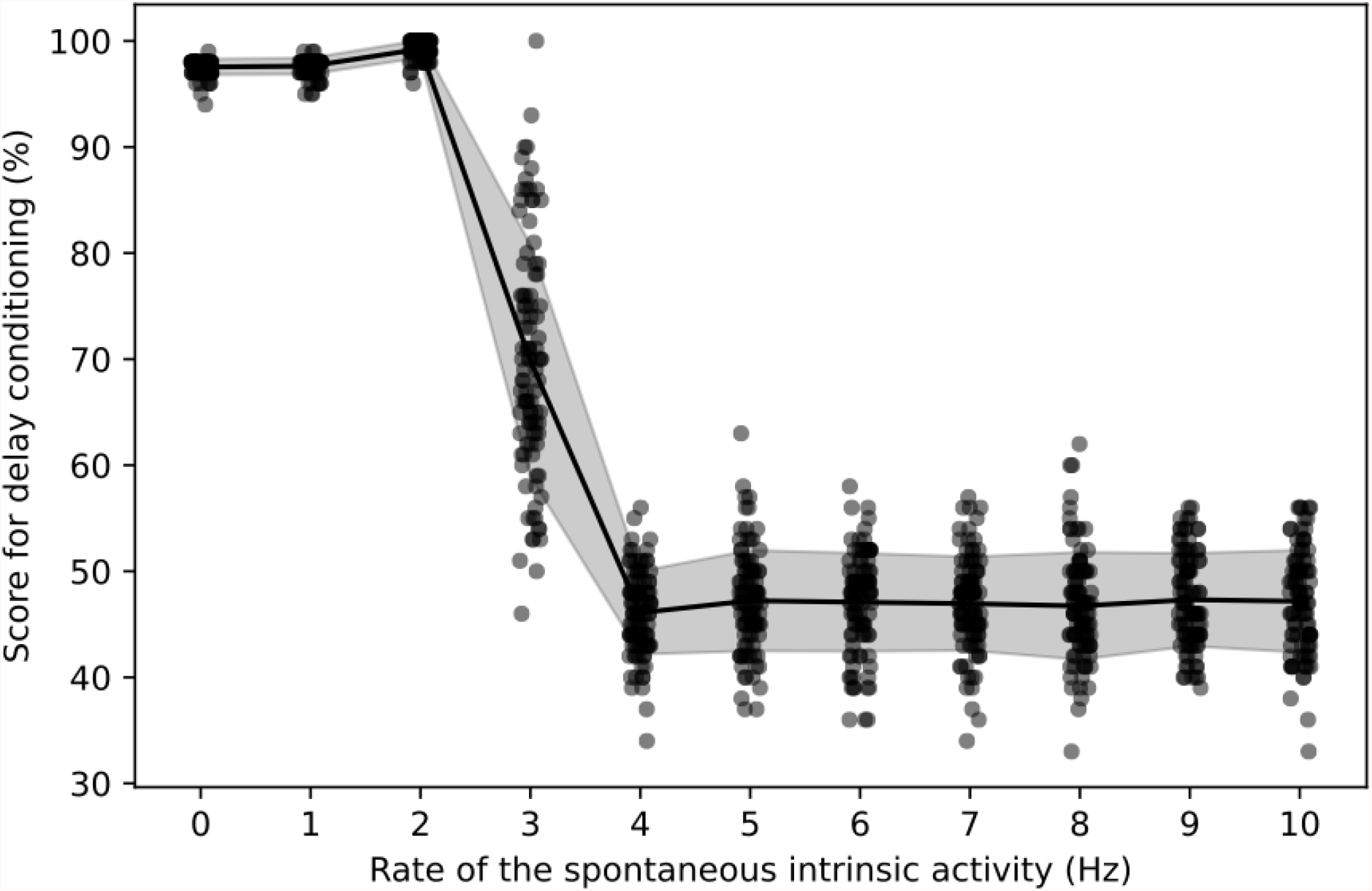
Effect of spontaneous intrinsic activity on the performance during the delay conditioning on the cognitive level. The accuracy of the binary classification as a function of the rate of spontaneous intrinsic activity. High values of spontaneous intrinsic activity force neurons to fire continuously, in an “epileptic” manner, and the network loses the ability to perform classification tasks. In such a state the average accuracy is slightly lower than a chance level (50%) because the Visual Cortex does not have 100% accuracy in digit recognition (Fig. 3d).

### 3. Conscious processing level - trace conditioning task

#### The critical role of inhibitory neurons

To date, the models associated with the Global Neuronal Workspace theory have focused on purely excitatory networks. Here we show the critical role of inhibitory neurons for conscious tasks. Additionally, we demonstrate how the performance on those tasks depends on the interplay between the ratio of excitatory and inhibitory neurons and spontaneous intrinsic activity.

Although the long-range connectivity itself is enough to perform the delay conditioning, it cannot achieve the trace conditioning task by itself (which is represented in Fig. 6a). On this level, the role of interneurons is seen to be critical in the function of the Prefrontal Cortex for the maintenance of conscious representation. This is needed to create and hold stable representations of stimuli and erase them when needed. Our model comprises two selective excitatory populations and one interneuron population. Once mature, it can hold a stable representation and erase it upon the presentation of a different stimulus (Fig. 6b). In this case, global epigenesis takes more time, on the order of ∼150 shown images, because it requires the local epigenesis in the PFC to occur first (Fig. 6c). Therefore, the mechanism of learning trace conditioning is highly dependent on the ability of working memory to maintain a needed stimulus.

**Figure 6.**
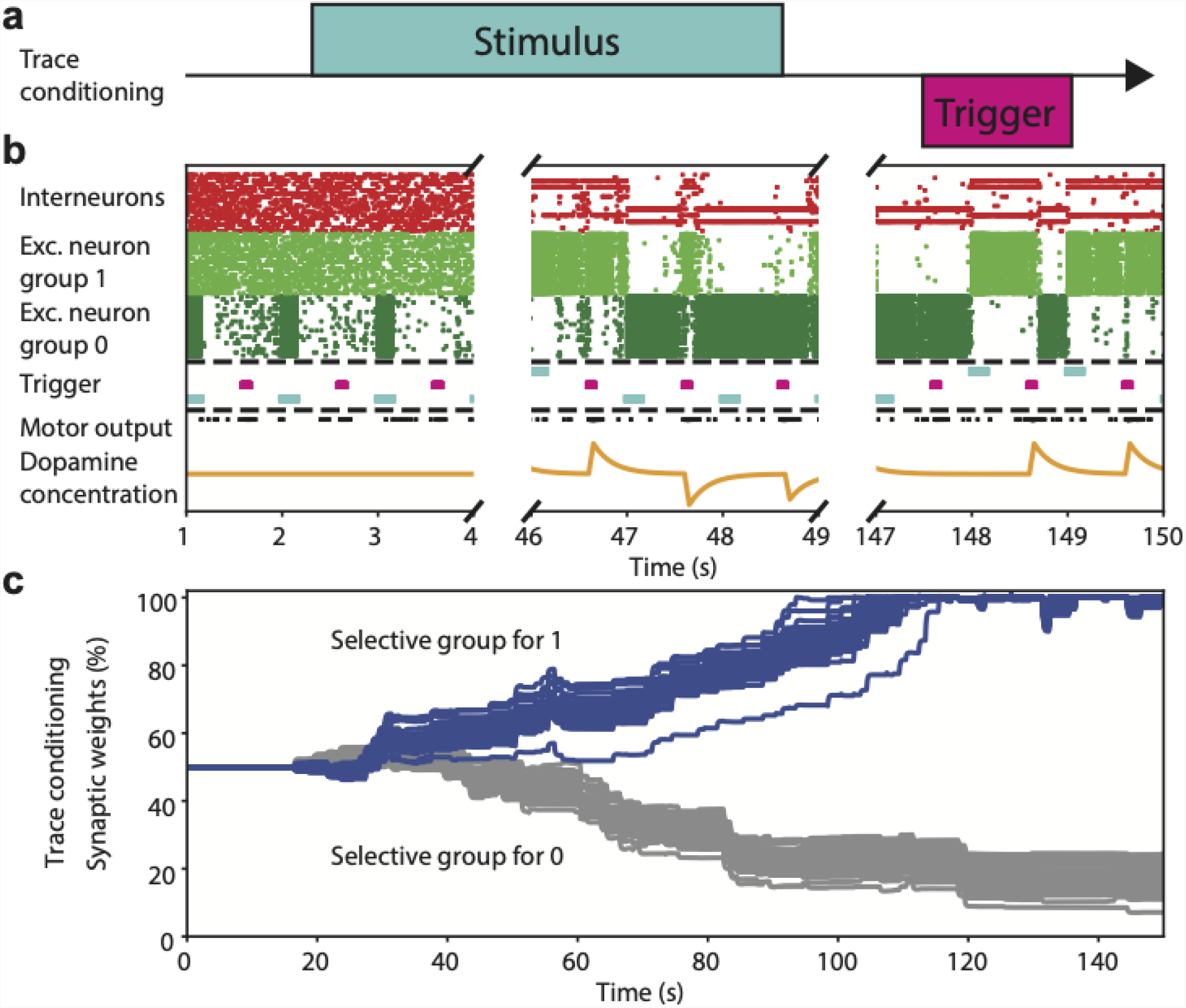
Epigenesis of the long-range connections network between Prefrontal and Motor Cortices while performing binary classification under trace conditioning. a. Description of the trace conditioning task. b. Evolution of the firing rate of the three populations of neurons within the GNW (inhibitory, excitatory selective for the digit 1, excitatory selective for the digit 0). 15 neurons per population are shown, each dot shows a neuron spiking. Depending on the network parameters, if the internal excitation in the GNW is not strong enough to cause constant feedback firing, the network gradually learns to sustain the representation of a stimulus in the GNW. When the next stimulus is present, the previous one is erased. The interneurons population also forms two distinct subpopulations selective to two different excitatory groups. c. Epigenesis of the global connections between Prefrontal and Motor Cortices. The network needs around 150 images to learn the task which is significantly more than with the delay conditioning because it also has to spend time learning how to hold representations in the GNW. Blue and grey lines correspond respectively to 1 and 0 output of the PFC. (describe the graphs 6b, 4b, (e.g. first 18 sec in 6b, or false negatives in 4b, mention the randomness of the dopamine and refractoriness, 2D graphs, learning time vs t(stim-trig), 6a timescale, and dop)

Just as in the intermediate level, with the delay conditioning we have studied how the model behaves with various values of the spontaneous intrinsic activity and the E/I ratio. The performance of the model strongly depends on both parameters, and high accuracy can be achieved only in the specific range of both values (Fig. 7a). In this case, we have identified a very specific range of spontaneous intrinsic activity (8-13 Hz), as well as a narrow interval of the E/I ratio which is located between 70% and 95% (maximum accuracy achieved with ratio = 80% and rate = 12Hz).

**Figure 7.**
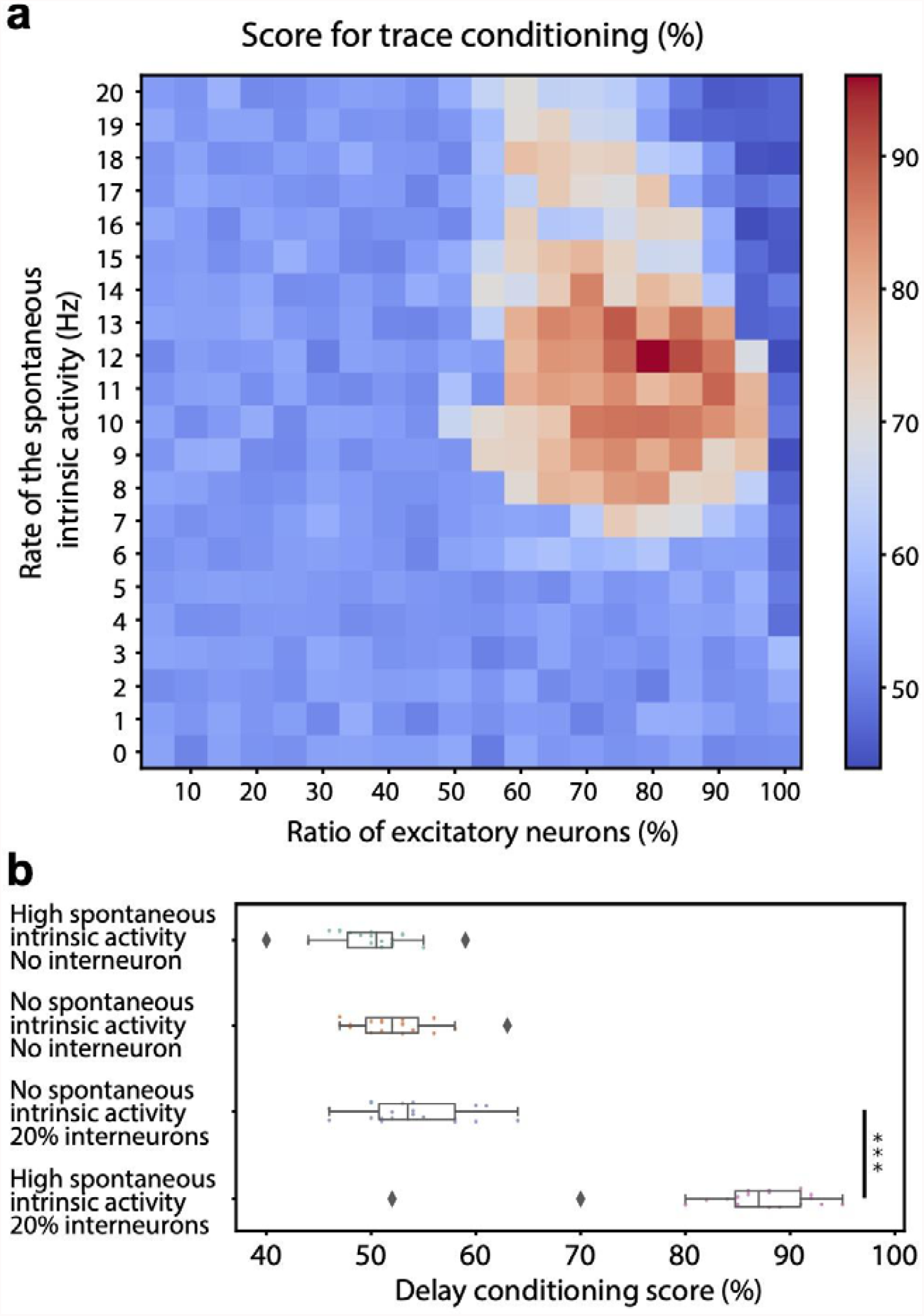
Factors affecting the performance during the trace conditioning on the Consciousness Level. **a**. The accuracy of the binary classification as a function of the rate of spontaneous intrinsic activity and the proportion of the interneurons. Unlike the delay conditioning on the cognitive level, the rate of the spontaneous activity plays a great role here, and in order for the model to show good performance (> 75%), both parameters must be within a specific range **b**. The accuracy of the model in four distinct states: 1) high spontaneous intrinsic activity and no interneurons; 2) no spontaneous intrinsic activity and no interneurons; 3) no spontaneous intrinsic activity and 20% of neurons are inhibitory; 4) high spontaneous intrinsic activity and 20% of neurons are inhibitory. The first three states provide poor performance, while the last one, which has the optimal set of parameters, yields high accuracy.

The same study has been conducted for the delay conditioning task to assure that the model can achieve good performance on the same parameter values (see Supplementary Figure 1). It is a crucial step towards biological plausibility: the network can not perform the trace conditioning task without being able to work with delay conditioning beforehand. In our case, we have identified the region where both tasks are performed with high accuracy.

## Discussion

In this paper, we address the issue of how the progressive complexification and integration of an artificial neural network gives rise to cognitive abilities. The current model is based on three hierarchical levels of information processing: the sensorimotor level, the cognitive level corresponding to the global nonconscious processing of information across different brain regions; and last the conscious level corresponding to the autonomous and lasting processing of information, even in the absence of externally applied sensory stimulation. This partition is consistent with the former distinction by Descartes and Kant of three levels of functional processing “Sensibility, Understanding and Reason” (see 31, 32). More recently, Daniel Kahneman proposed the dichotomy of a fast “System 1”, instinctive and emotional, and a slow “System 2”, which is more deliberative and logical (6). The development of computer science and artificial intelligence have led to a strikingly different distinction such as David Marr’s partition of computational mechanisms into what he refers to as “formal levels”: Implementation, Algorithms, and Computations (41). Marr’s claim relied on the underlying assumption that the computations required for a given cognitive task are distinct from the physical implementation, natural or artificial, of the network. They are independent of the cognitive process under study and most of all, of the level of organization of the network. The word “level” has in both cases entirely different meanings.

In light of the need for grounding cognitive models in biology and validating their genuineness across levels, we followed successive mechanisms in known structural and functional organizations, identified through both the development and evolution of the human brain (42). The first step proceeds from local to global integration of information; a second one moves from nonconscious to conscious processing. This leads to the three structurally and physiologically grounded levels: “perception,” with the learning of invariants through local synaptic epigenesis, “cognition” with the learning of nonconscious integration through the long-range connectivity and reward, and finally, conscious processing with the learning of maintenance of representation online through interneurons and spontaneous activity. The model expresses key insights across the three levels of analyses: first, synaptic epigenesis, modeled here by selection and stabilization of synapses, is a critical mechanism at all levels, from perception to consciousness; second, dopamine is necessary for cognitive tasks to achieve proper credit assignment despite temporal delay between perception and reward; thirdly, interneurons allow the maintenance of self-sustained representation within the GNW in the absence of sensory input, thus enabling the system to solve conscious tasks. Finally, our results show how balanced spontaneous intrinsic activity facilitated epigenesis at both local and global scales, the balanced excitatory-inhibitory ratio increased performance. Those observations, in addition to emphasizing general principles at play in the human brain, are in surprising accordance with empirical observations. For instance, without including synaptic pruning, the temporal evolution of synaptic weights leads to the three classical phases of growth, maximal variability, and stabilization (38). More unexpectedly, the optimal neurobiological parameters predicted for the conscious level are 20% of inhibitory neurons, which is predicted for balanced networks (43). Lastly, the “optimal” 12 Hz of spontaneous activity discovered in our artificial network coincides with the actual recordings of spontaneous spiking in the human cortex and is also strongly reminiscent of the highly conserved and omnipresent alpha rhythm in the cerebral cortex (44).

Computational neuroscience has increasingly relied on multi-scale brain models (45, 46) to explain basic cognitive functions (47), active, top-down, and prospective memory retrieval (48, 49), and even syntactic processing for the production of language (50). Some of these models, like ours, have employed spiking neural networks with STDP to study synaptic epigenesis during learning of sensory representation for tactile (51) and visual (37) perception, or to demonstrate how biases in natural statistics can influence population encoding and downstream correlates of behavior (52). These models, however, have tended to focus on a single learning mechanism, applied to a specific task. Here we combine both Hebbian and reinforcement learning across different canonical tasks. While the tasks considered here may appear minimalistic compared to recent sensational breakthroughs coming from artificial intelligence (53, 54) our goal was not to achieve behavioral complexity but to uncover general biological principles through natural plausibility. Moreover, while it is certainly possible for artificial systems to mimic the behavior of a conscious agent, it is far less trivial to demonstrate the genuineness of its semantic understanding and subjective experience (55). From a purely pragmatic viewpoint, the Turing test (56) relies only on the judges being unable to tell if the agent is a human or a machine (57). We chose to view the problem in another way, by grounding the cognitive architecture in neurobiology and from there building the most parsimonious, realistic model capable of solving both perceptual and conscious tasks. This allowed delineating necessary and sufficient biological mechanisms for cognitive abilities in an artificial neural network (58). Future works should probe the role of other important biological mechanisms such as oscillations (59) or distributed coding for task value (60), and may likewise wish to take into account additional subcortical structures (beyond the striatum), such as the thalamus (61) and the basal ganglia (62). A promising avenue is also to explore the combination of the GNW architecture with predictive processing (63); the top-down feedback required for ignition and conscious access then becomes entangled with the prediction signal of the internal generative model of the world (64). Finally, the GNW architecture is also opening new venues in deep learning: on the one hand, by creating high-level inductive biases that improve out-of-distribution generalization (65, 66), and, on the other, by providing an amodal latent space where alignment of representations across modalities become automatic (67).

Recently, a separate distinction was made between basic “conscious’ processing and “the self-monitoring of those computations, leading to a subjective sense of certainty or error” (68). This metacognitive dimension of consciousness is also linked to the hypothesis of the social origin of consciousness, which proposes the sense of self as a vestige of the evolutionary skills initially developed for understanding others (69). In this sense, the hominization of primates seems connected to molecular changes associated with social cognition (70) and, in line with our results, with the increase of long-range connectivity and regulation of the balance between excitatory and inhibitory neurons (71). Beyond these features, the prolonged postnatal brain development with a proper cascade of critical periods (72, 73) leverages the multiple non-genetic interactions with the physical, social, and cultural environment, ultimately, giving rise to categorically human-specific cognitive abilities including the recursivity of language (71, 74, 75). A key perspective is thus to further investigate the role of cultural embedding, adding explicitly a social dimension to the global neuronal workspace (17, 76). Some dyadic computational models (77, 78) have already been proposed using either continuous dynamical systems or discrete symbolic representation. Anchoring future computational models in both biological and social realities will not only continue to shed light on the core mechanisms underlying cognition, but it will also help to provide a unique bridge to artificial intelligence towards the only known systems with advanced social consciousness: the human brain (79).

## Materials and Methods

### Model

#### 1. Local Network

##### S1 layer

The convolution of the images is done with six 7×7 Gabor filters. The frequency is 0.1 and the standard deviation is 1 for all of them. The angles of the filters start at π/8 and go with the step of π/6. The offset of π/8 was introduced in order not to focus on vertical and horizontal lines which are not as informative in classification. Every time the Gabor kernel overlaps a border of a picture, this picture is extended by replicating the edge pixels. The resulting convolution is again resized to 128×128 pixels, thus creating an S1 layer with the size of 128×128×6.

##### C1 layer

Each cell of this layer processes spikes from S1 cells in a 7×7 square (receptive field size of a given C1 cell). A C1 cell propagates only the single strongest orientation from its receptive field at a time. The receptive fields of two adjacent C1 cells are shifted by 6 pixels, meaning that they have an area of intersection of 7×1 pixels. This leads to a total C1 layer size of 25×25×2 (25×25 for the propagated values and 25×25 to store the orientations). However, the C1-S2 connections need to be set in a way that helps to identify not only the intensity of the propagated signal but the orientation. To implement this in a more biologically plausible way, the C1 layer has to be transformed. Each element on the 25×25 grid would now contain 6 C1 cells (one for each orientation) with the winner-takes-all condition: only one out of 6 cells can fire at a given time. Now, if we want to trace the signal back we can determine not only its spatial location but also its orientation. This modification leaves the C1 layer with a size of 25×25×6 with only 25×25 cells firing each time. We have also implemented the lateral inhibition procedure in a way it is done in the Masquelier and Thorpe model (37). The C1-S2 connections were governed by STDP.

##### S2 layer

The S2 layer is represented by a fixed number of integrate-and-fire neurons (we used either 10 or 20 neurons). We have also added a sinusoidal function into the equations of S2 cells to model the alpha waves. By doing this we achieve a clearing effect of periodic modulations. The wave and the images are not perfectly in phase with each other and the modulation wave is shifted by an angle of π/8. This provides better elimination of the spontaneous intrinsic activity and signal amplification. To determine which S2 cells encode which numbers the mature network was presented with a test set of images. To perform such classification the winner-takes-all strategy is used once more: a stimulus gets propagated to only one cell which fires first. Then each S2 cell was assigned to a number to which it fired the greatest number of times.

#### 5. Full Network

##### Prefrontal Cortex

The population is represented by 100 integrate-and-fire neurons. 80 neurons are excitatory and they are connected to the S2 layer of the Visual Cortex network. The excitatory population is divided into selective groups in a way that the number of groups is equal to the number of digits presented in the input (e.g., 2 groups if we use only images of 0 and 1). Each selective group is connected to S2 patterns responsible for one particular digit. The other 20 are interneurons and they play an inhibitory role instead. The connection pattern in the Prefrontal Cortex is all-to-all.

Such a state of the Prefrontal Cortex model was used to perform trace conditioning. To create the intermediate level for performing nonconscious delay conditioning, all the interneurons were deleted from the model.

##### Motor Cortex

The model of the Motor Cortex uses only one integrate-and-fire neuron. It gets input from the excitatory population of the Prefrontal Cortex. Its output is considered to be a sign of transmission of the information to the external environment and it induces the activation of the reward pathway.

##### Striatum

To perform the reinforcement learning we introduce one integrate-and-fire neuron responsible for changing the extracellular dopamine concentration in the Prefrontal Cortex. It can be seen as a model of the Striatum. The dopamine concentration is set on a baseline and increases or decreases depending on whether the answer was right or wrong. Before the next image is presented the concentration exponentially decays back to the baseline.

#### 6. STDP mechanisms

In our model, there are three areas where synaptic populations are subject to epigenesis: C1-S2 connections in the local network; PFC-Motor Cortex, and PFC-PFC connections (inhibitory and excitatory) in the full network. We use different models of STDP to set up different connections.

##### Classical STDP

This is the main plasticity model in our networks that is used in all excitatory connections except PFC-Motor Cortex. We used the following equations to model the effect of STDP:

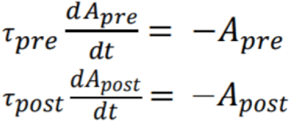

This system of equations models two possible scenarios: the firing of a presynaptic neuron and the firing of a postsynaptic neuron. In the first case, the value of A_pre_ is updated by adding an increment value ΔA_pre_ > 0 (models the effect of further potentiation), and the synaptic weight is updated by A_post_ in a given moment of time (depression). In the second case, everything is the opposite: A_post_ is updated by adding an increment value ΔA_post_ < 0 for future depression, and the weight is potentiated by the value of A_pre_.

##### Symmetrical STDP

The symmetrical model of STDP is based on the STDP measured experimentally in the GABAergic neurons population (80). It is used to govern the plasticity of interneurons in the Prefrontal cortex. The mechanisms for the symmetrical STDP are similar to the classical STDP however the equations are slightly different:

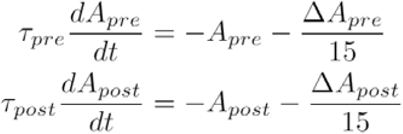

where ΔA_pre_ > 0 and ΔA_post_ > 0. The additional terms are added to keep the values of A_pre_ and A_post_ slightly below zero to model the depression when two neurons fire with a large temporal difference. In other cases, when the temporal distance is short enough the model produces an effect of the long-term potentiation.

##### Dopamine modulated STDP

This type of synaptic plasticity is used to introduce reinforcement learning which involves multiple brain regions and feedback from the external environment. The model is based on the work of Izhikevich on linking STDP and dopamine signaling (33). It is

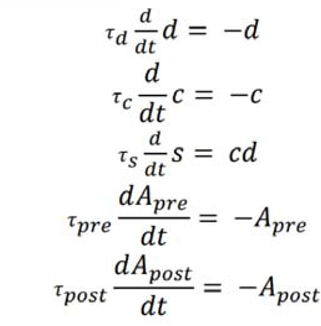

represented by a following set of equations:

The variable *‘c’* is updated in the same way as the synaptic weight is updated during classical STDP (Using equations 4 and 5). The variable *‘d’* represents the extracellular dopamine concentration: if it is lower than a baseline, then it launches the negative reward mechanism, if higher - positive. The variable ‘*s’* serves as an increment to the synaptic weight and depends not only on the sign and magnitude of the classical STDP but on the sign and magnitude of the variable *‘d’*. The release of dopamine is controlled by a single neuron that fires once when a new image is presented with a random delay from 0 to 1/32 seconds.

## Acknowledgements

J.P.C. has received funding from the European Union’s Horizon 2020 Framework Programme for Research and Innovation under Specific Grant Agreement No. 945539 (Human Brain Project SGA3). G.D. is funded by the Institute for Data Valorization (IVADO; CF00137433), Montreal, and the Fonds de recherche du Québec (FRQ; 285289). Authors are grateful for the constructive comments of Samuel Bollota, David Yu-Tung Hui, and Timothy Nest.

## Supplementary Material

**Supplementary Figure 1.**
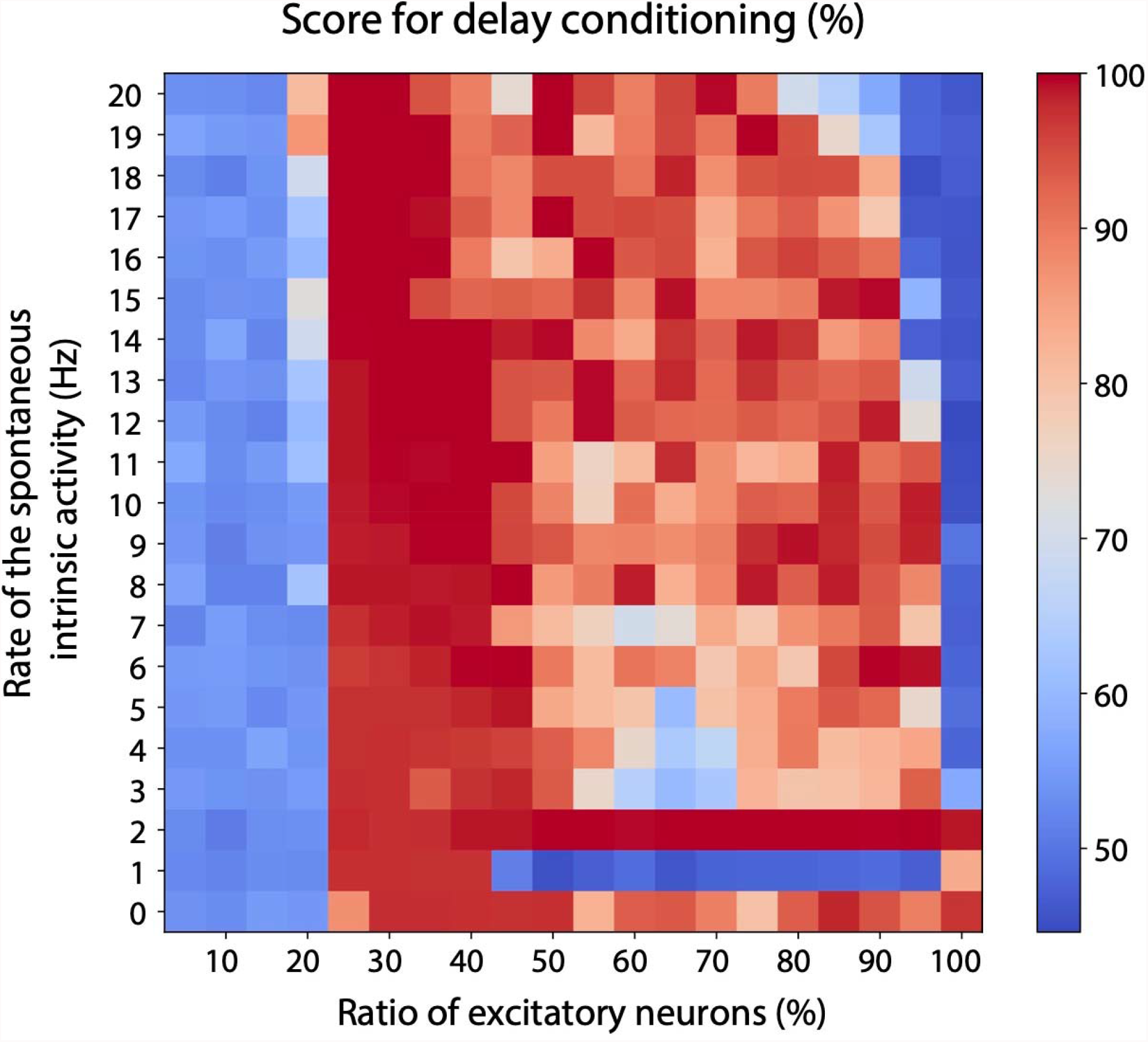
Factors affecting the performance during the delay conditioning on the *cognitive level*. The accuracy of the binary classification as a function of the rate of spontaneous intrinsic activity and the proportion of the interneurons. Notice how the optimal parameters for the trace conditioning on the Conscious Level still provide good performance for the delay conditioning.

